# Dynamic recycling of extracellular ATP in human epithelial intestinal cells

**DOI:** 10.1101/2023.02.10.527987

**Authors:** Nicolas Andres Saffioti, Cora Lilia Alvarez, Zaher Bazzi, María Virginia Gentilini, Gabriel Gondolesi, Pablo Julio Schwarzbaum, Julieta Schachter

**Affiliations:** Instituto de Química y Físico-Química Biológicas “Prof. Alejandro C. Paladini”, Universidad de Buenos Aires (UBA), Consejo Nacional de Investigaciones Científicas y Técnicas (CONICET), Facultad de Farmacia y Bioquímica, Junín 956, Buenos Aires, Argentina; Universidad de Buenos Aires (UBA), Facultad de Farmacia y Bioquímica, Departamento de Química Biológica, Cátedra de Química Biológica, Junín 956, Buenos Aires, Argentina; Instituto de Nanosistemas, Universidad Nacional de General San Martin, Argentina; Fundación Favaloro Hospital Universitario, Unidad de Insuficiencia, Rehabilitación y Trasplante Intestinal, Buenos Aires, Argentina; Instituto de Medicina Traslacional, Trasplante y Bioingeniería (IMETTyB, CONICET, Universidad Favaloro), Laboratorio de Inmunología asociada al Trasplante, Buenos Aires, Argentina; Universidad de Buenos Aires (UBA), Facultad de Ciencias Exactas y Naturales, Departamento de Biodiversidad y Biología Experimental, Intendente Güiraldes 2160, Ciudad Universitaria, Buenos Aires, Argentina

## Abstract

Intestinal epithelial cells play important roles in the absorption of nutrients, secretion of electrolytes and food digestion. The function of these cells is strongly influenced by purinergic signalling activated by extracellular ATP (eATP) and other nucleotides. The activity of several ecto-enzymes determines the dynamic regulation of eATP. In pathological contexts, eATP may act as a danger signal controlling a variety of purinergic responses aimed at defending the organism from pathogens present in the intestinal lumen.

In this study, we characterized the dynamics of eATP on polarised and non-polarised Caco-2 cells. eATP was quantified by luminometry using the luciferin-luciferase reaction. Results show that non-polarized Caco-2 cells triggered a strong but transient release of intracellular ATP after hypotonic stimuli, leading to low micromolar eATP accumulation. Subsequent eATP hydrolysis mainly determined eATP decay, though this effect could be counterbalanced by eATP synthesis by ecto-kinases kinetically characterized in this study. In polarized Caco-2 cells, eATP showed a faster turnover at the apical vs the basolateral side.

To quantify the extent to which different processes contribute to eATP regulation, we created a data-driven mathematical model of the metabolism of extracellular nucleotides. Model simulations showed that eATP recycling by ecto-AK is more efficient a low micromolar eADP concentrations and is favored by the low eADPase activity of Caco-2 cells. Simulations also indicated that a transient eATP increase could be observed upon the addition of non-adenine nucleotides due the high ecto-NDPK activity in these cells. Model parameters showed that ecto-kinases are asymmetrically distributed upon polarization, with the apical side having activity levels generally greater in comparison with the basolateral side or the non-polarized cells.

Finally, experiments using human intestinal epithelial cells confirmed the presence of functional ecto-kinases promoting eATP synthesis. The adaptive value of eATP regulation and purinergic signalling in the intestine is discussed.

**Authors summary:** Intestinal epithelial cells play important roles in the absorption of nutrients, secretion of electrolytes and food digestion. When intracellular ATP is released into the intestinal milieu, either at the lumen or the internal side, the resulting extracellular ATP can act as an alert signal to engage cell surface purinergic receptors that activate the immune defence of the organism against pathogens.

We worked with Caco-2 and primary human intestinal cell, and our results showed that extracellular ATP regulation is a complex network of reactions that simultaneously consume or generate ATP in whole viable intestinal epithelial cells. In particular, we created a mathematical model, fitted to experimental data, that allowed to quantify the degree to which intracellular ATP release and the activity of a variety of ectoenzymes controlling the concentration of extracellular ATP in a complex way.

## 1. Introduction

The surface of the intestine is covered by a layer of cells that form the intestinal epithelium. Intestinal epithelial cells play important roles in the absorption of nutrients, secretion of electrolytes, digestion of food and host defence mechanisms [1, 2]. The function of intestinal epithelial cells is strongly influenced by extracellular nucleotides, supporting a complex signalling network that mediates short-term functions such as secretion and motility, and long-term functions like proliferation and apoptosis [3, 4]. Among these nucleotides, extracellular ATP (eATP) was found to be an early danger signal response to infection with enteric pathogens that eventually promote inflammation of the gut [4, 5].

An important source of eATP is the intracellular ATP (iATP) found in the cytosol and vesicles of many cell types [6]. Activation of iATP release was found in subepithelial intestinal fibroblasts, human epithelial cell lines and enteroendocrine cells in response to several stimuli, including agents that elevate cAMP, such as forskolin and cholera toxin [7], low medium phosphate, hypoosmotic swelling and bacterial infection [7, 8]. Currently, several ATP conduits have been postulated to mediate regulated iATP release, including various anion channels, connexins and pannexin-1 hemichannels and the calcium homeostasis modulator 1 [6].

Extracellular ATP and other di- and tri-phosphonucleosides can activate purinergic receptors 2 (P2 receptors) unevenly distributed in the small and large intestine [9]. Purinergic signalling is controlled by membrane bound ecto-nucleotidases and ecto-kinases capable of promoting the synthesis and/or hydrolysis of eATP, and/or its conversion into other extracellular nucleotides and nucleosides. For any cell type and metabolic context, a specific set of ecto-enzymes may control the rate, amount and timing of nucleotide turnover [10].

Ecto-nucleoside triphosphate diphosphohydrolases (Ecto-NTPDases) are a family of enzymes promoting the extracellular hydrolysis of eATP, eADP, eUTP and eUDP. One or more members of this family are present in almost every cell. Ecto-NTPDase-1, -2, and -3, which differ regarding the specific preferences for nucleotides, are responsible for the hydrolysis of nucleoside diphosphates (NDPs) and nucleoside triphosphates (NTPs) in various tissues of the gastrointestinal tract [1]. Regarding eATP and eADP hydrolysis, ecto-NTPDase-1 hydrolyses both nucleotides at similar rates, while ecto-NTPDase-2 has a high preference for eATP over eADP and ecto-NTPDase3 is a functional intermediate which preferably hydrolyses eATP [11].

The intestinal cell line HT29 cells expressed functional ecto-NTPDase-2 displaying high ecto-ATPase activity [12], while Caco-2 cells and their exosomes were reported to exhibit ecto-NTPDases-1 and -2 at the cell membrane [13, 14].

Extracellular ATP can be also metabolized by ecto-kinases, with ecto-adenylate kinase (Ecto-AK) facilitating the reversible conversion of eADP to eATP and eAMP, and ecto-nucleoside diphosphate kinase (Ecto-NDPK) promoting the exchange of terminal phosphate between extracellular NDPs and NTPs [10]. All these ecto-enzymes, if present and active, should be able to control the concentration of eATP.

Up to now, although some ecto-enzymes have been identified in intestinal cells, no attempts have been made to characterize the dynamic interaction of these membrane proteins on eATP regulation of intestinal cells. In this study, we aimed to characterize iATP release and eATP recycling by ecto-enzymes, contributing to eATP regulation in Caco-2 cell line. The Caco-2 cells derive from colorectal adenocarcinoma and easily differentiate into cells exhibiting the morphology and function of enterocytes, the absorptive cells of the small intestine [15]. The experimental studies on eATP dynamics in polarized and non-polarized Caco-2 were complemented with a mathematical model quantifying the complex relationship among the different processes contributing to eATP regulation. Our results provide a quantitative description of the eATP dynamics of human intestinal epithelial cells.

## 2. Results

In this section we show experimental results on eATP kinetics of non-polarized and polarized Caco-2 cells. To understand the dynamics of the different processes contributing to eATP regulation, a mathematical model was fitted to experimental data, and predictions were made. Finally, for a comparative purpose, we show results of a few experiments made on epithelial cells obtained from intestinal surgical pieces.

### 2.1. Non-polarized Caco-2 cells

#### 2.1.1. eATP kinetics after hypotonic shock

The kinetics of eATP accumulation, *i.e.*, eATP kinetics, results from the dynamic balance between iATP release mechanisms and the activities of ecto-enzymes capable of degrading and/or synthetizing eATP. As a first step towards the characterization of eATP kinetics, iATP release was triggered by exposing Caco-2 cells to hypotonic media (Fig 1).

**Figure 1.**
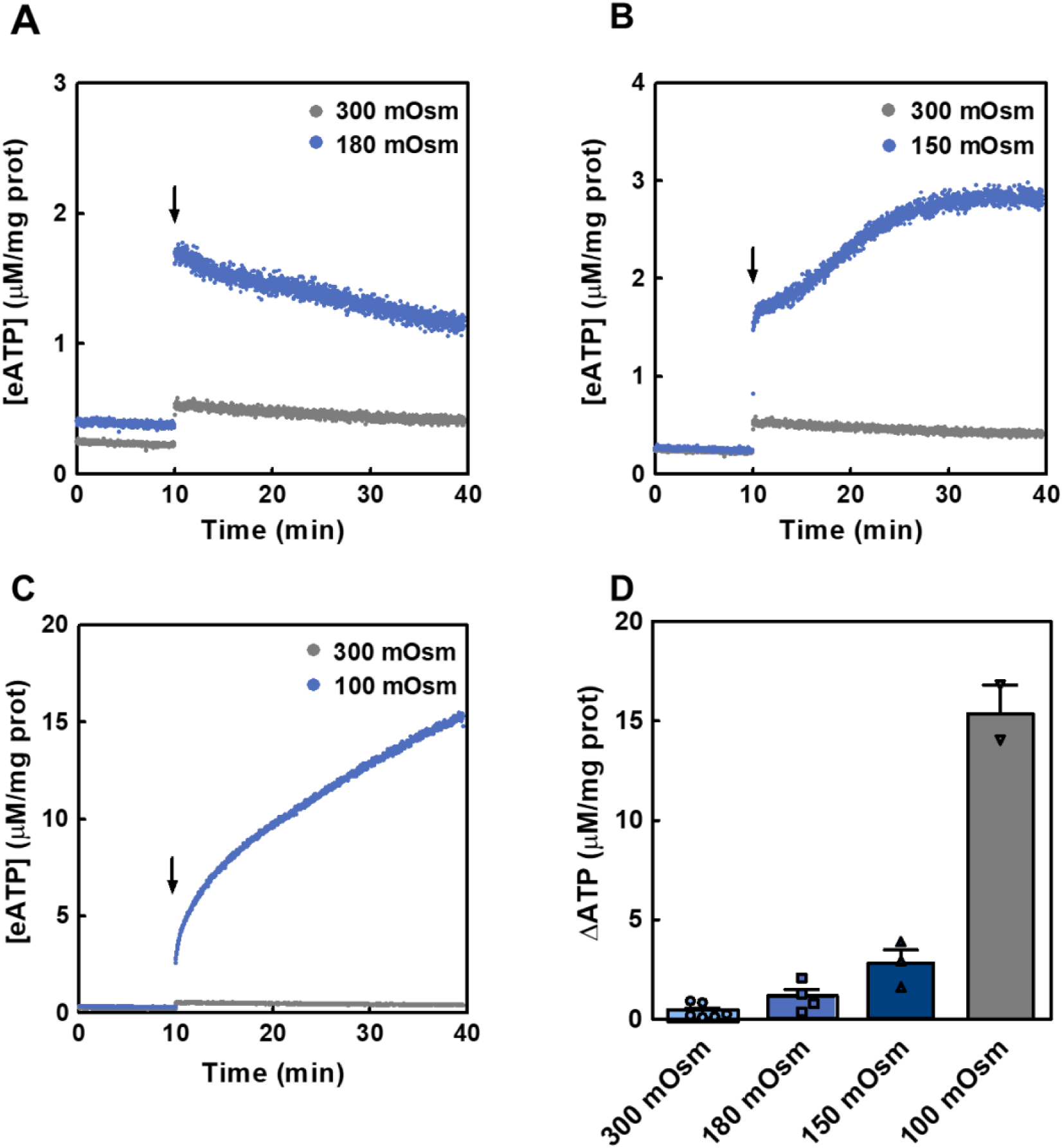
eATP kinetics of hypotonically-stimulated Caco-2 cells. The time course of [eATP] from Caco-2 cells after a hypotonic shock was quantified by luminometry performed at room temperature. (A-C) Cells were maintained in isotonic medium and, at the times indicated by the arrow, were exposed to isotonic medium (grey) or to hypotonic media (blue) of 180 mOsm (A), 150 mOsm (B) and 100 mOsm (C). Results are expressed as means of [eATP] from 4, 3 and 2 independent experiments run in triplicate for the 180, 150 a 100 mOsm experiments, respectively. (D) Increases in [eATP] from data in A-C were evaluated as ΔATP, i.e., the difference between [eATP] at 30 minutes post-stimulus and basal [eATP]. Cells were exposed to 300 mOsm (light blue bars), 180 mOsm (blue bars), 150 mOsm (dark blue bars) and 100 mOsm (grey bars). Bars show mean values + standard error of the mean (s.e.m) from 2 to 5 independent experiments. Points represent the independents values for each condition.

Under unstimulated conditions, [eATP] remained stable. Whereas addition of isotonic medium triggered a slight increase of [eATP], hypotonic media (100-180 mOsm) activated a stronger iATP release with different kinetics according to the osmotic gradient imposed (Fig 1A-C). As shown in Fig 1D, [eATP] increased non-linearly with the magnitude of the hypotonic stimulus.

The experimental [iATP] amounted to 1.81 mM. By comparing [iATP] with [eATP] along eATP kinetics, it was possible to estimate the energy cost of iATP release. Calculations were made for cells exposed to isotonic or 180 mOsm media, two conditions where no lysis was detected [14]. During the isotonic shock, representing a mechanical stimulus in the absence of osmotic gradient, eATP amounted to 0.33% of iATP, while under 180 mOsm this figure amounted to 3.6%. Thus, the energy cost of eATP production by iATP efflux was very small (see section 4.9 for further details). No iADP release was detected in the 180 mOsm stimulus (S1 Fig)

In our previous work, we showed that ecto-nucleotidases present in Caco-2 cells catalyse significant rates of eATP hydrolysis, leading to eADP accumulation [14]. In principle, the resulting accumulated eADP could be used by the potential presence of ecto-kinases like ecto-AK and ecto-NDPK, present in several cell types, to synthetize eATP. Thus, in the following experiments the activities of ecto-AK and ecto-NDPK were assessed by quantifying eATP kinetics under different conditions.

#### 2.1.2. Ecto-AK activity in Caco-2 cells

AK catalyses the following reversible reaction: 2 eADP ↔ eATP + eAMP and is inhibited by Ap5A [16]. Ecto-AK activity was then assessed by following eATP synthesis when Caco-2 cells were incubated with exogenous eADP (6-48 µM). Non-linear [eATP] increases were proportional to [eADP] (Fig 2A). At 30 minutes post-stimulus, treatment with 10 μM Ap5A, which does not permeate cells, inhibited eATP synthesis by 100 % (6-24 µM eADP) or by 90% (48 µM eADP) (Fig 2B), thus showing the presence of a functional ecto-AK in Caco-2 cells membrane.

**Figure 2.**
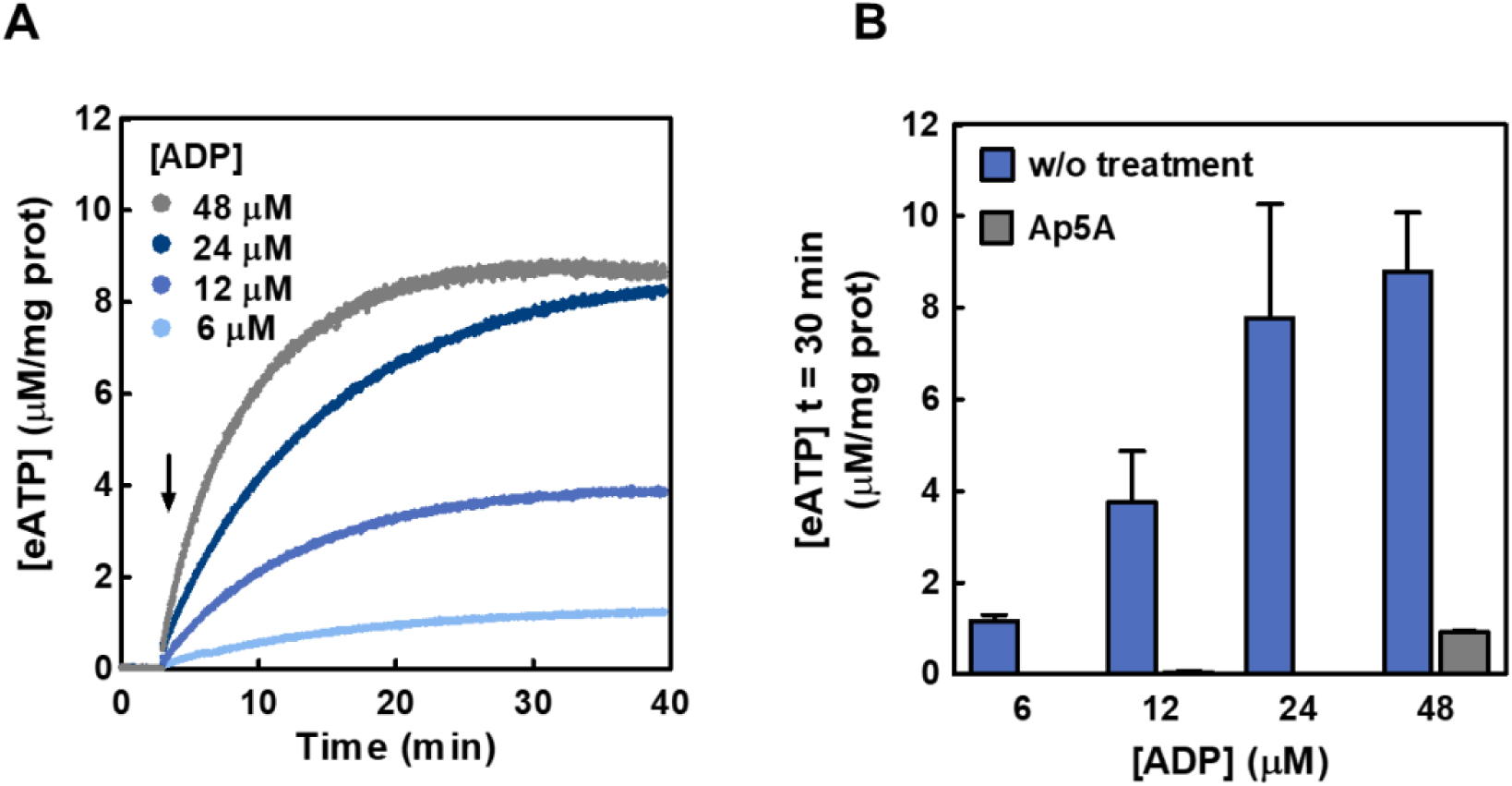
Synthesis of eATP from eADP in Caco-2 cells. (A) The time course of [eATP] synthetized from exogenous eADP (6-48 μM) in the extracellular medium of intact Caco-2 cells was quantified by luminometry. The cells were incubated with the luciferin - luciferase reaction mix and the [eADP] indicated in the figure were added at the time indicated by the arrow. Data are means of at least 3 independent experiments run i n duplicate for each [eADP]. (B) Effect of treatment with Ap5A (adenylate kinase inhibitor) on eATP synthesis from eADP in Caco-2 cells. The cells were treated or not (w/o treatment) with 10 μM Ap5A and the [eATP] at 30 minutes was measured by luminometry under similar conditions as experiments in (A). The bars are means ± s.e.m. from at 3-5 independent experiments run in duplicate in the absence of Ap5A and 2 independent experiments in the presence of the inhibitor.

#### 2.1.3. Ecto-NDPK activity in Caco-2 cells

NDPK catalyses the transfer of a γ–phosphate from NTP to NDP. Thus, in the presence of eADP and a given eNTP, the following reaction: eADP + eNTP ↔ eATP + eNDP leads to eATP synthesis when eADP is phosphorylated by NDPK.

Accordingly, incubation of cells with 100 μM eCTP at different [eADP] (3-12 μM) resulted in the rapid synthesis of eATP (Fig 3A). Maximal [eATP] values were obtained 30 minutes after the addition of substrates (Fig 3A). The experiments were conducted in the presence of 10 μM Ap5A to rule out any contribution of ecto-AK to the observed eATP kinetics.

**Figure 3.**
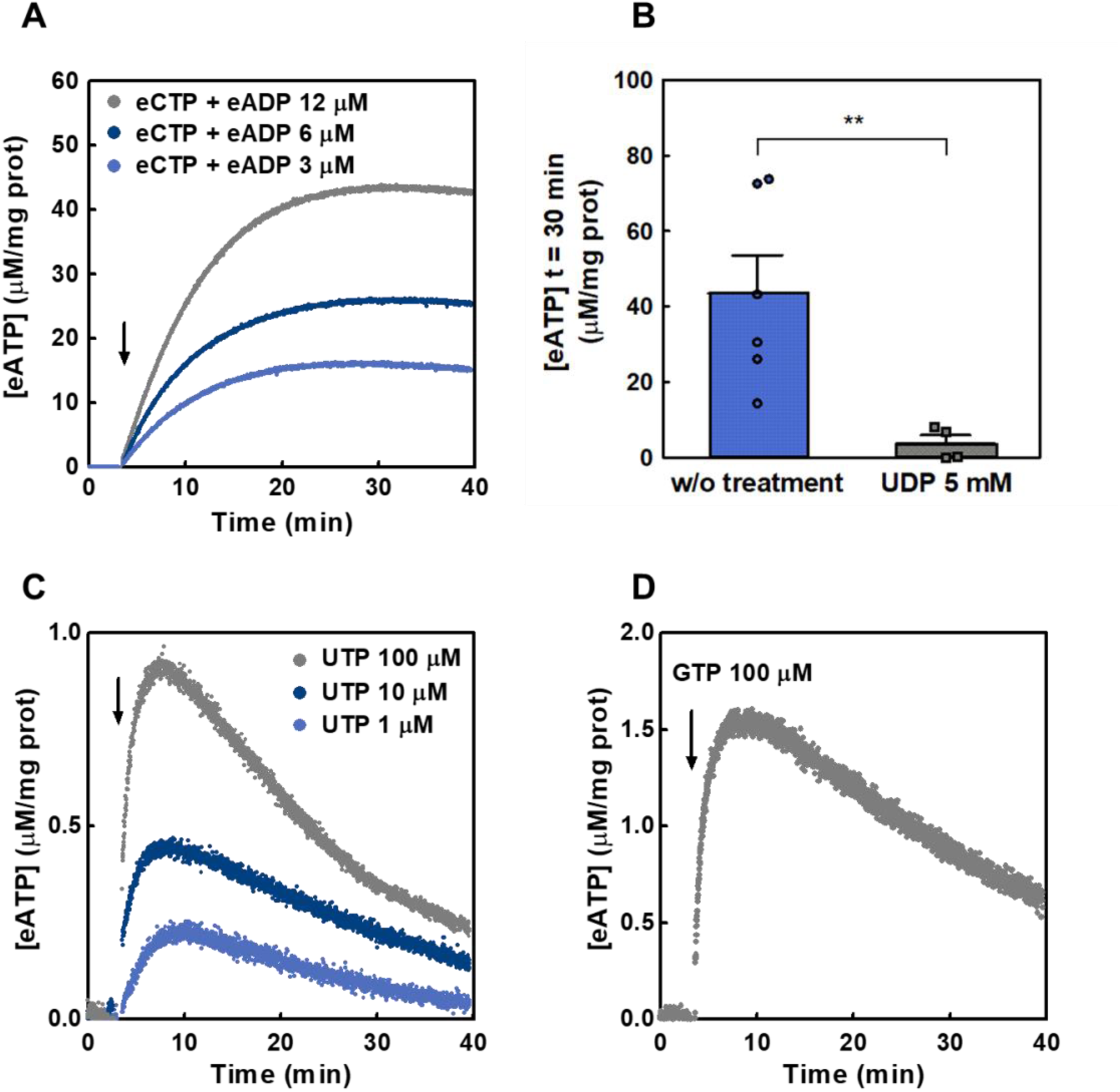
Extracellular synthesis of eATP from eADP and eCTP, eUTP and eGTP in Caco-2 cells. (A) The time course of eATP synthesis from eCTP (100 μM) and eADP (light blue for 3 μM, blue for 6 μM and grey for 12 μM) in the extracellular medium of Caco-2 cells was quantified by luminometry. Cells were incubated with the reaction mix and eCTP and eADP were added at the time indicated by the arrow. Data are means of 3 to 5 independent experiments run in duplicate for each ADP concentration. (B) Production of eATP after 30 minutes exposure of Caco-2 cells to 100 μM eCTP and 12 μM eADP. Experiments were run in the presence of 5 mM eUDP (grey bar and squares) or in its absence (blue bar and points). The [eATP] was measured under conditions similar to the experiments in (A). Results are expressed as [eATP] in μM/mg of protein, bars are means ± s.e.m from 4 to 7 independent experiments run in duplicate. ** means P-value <0.01 in comparison with the condition without treatment. (C) and (D) The time course of eATP accumulation in the presence of eUTP (C; grey for 100 μM, dark blue for 10 μM, blue for 1 μM) or 100 µM eGTP (D). Data are the means from 4 independent experiments in the case of 100 µM eUTP, 3 in the case of 100 eGTP, µM and 2 independent experiments in the case of 10 or 1 µM eUTP. Nucleotides were added at the time indicated by the arrow.

Addition of 5 mM eUDP, together with 100 μM eCTP and 12 μM eADP, decreased the eATP synthesis by 91% (Fig 3B), a result compatible with high [eUDP] favouring eUDP to eUTP conversion by ecto-NDPK, rather than eATP synthesis from eADP.

In separate experiments, addition of increasing [eUTP] (1-100 μM) without the addition of exogenous eADP (only endogenous eADP present), resulted in a concentration-dependent increase of [eATP] (Fig 3C). Because this increase was abolished by 5 mM eUDP (S2 Fig), we hypothesized that eATP synthesis was due to ecto-NDPK activity using exogenous eUTP and endogenous eADP. This is because there is a basal eADP concentration in the extracellular media of 0.77 ± 0.47 μM eADP/mg protein (S3 Fig). A similar experiment using 100 µM eGTP, instead of eUTP, provided qualitatively similar results (Fig 3D). Overall results showed a functional ecto-NDPK activity capable of synthetizing eATP from different γ-phosphate donors (eCTP, eUTP and eGTP) in the presence of endogenous and exogenous eADP.

#### 2.1.4. Modelling eATP kinetics of non-polarized Caco-2 cells

Caco-2 cells regulate eATP kinetics by iATP release, eATP synthesis by the activities of ecto-AK and ecto-NDPK (as shown in this study), and hydrolysis by ecto-nucleotidases [14]. Thus, to quantify the contribution of these processes to eATP kinetics, we built a mathematical model that was then fitted to experimental data.

A scheme of the model is depicted in Fig 4A. In the model, [eATP] can increase by iATP release, by lytic and by non-lytic mechanisms. In addition, [eATP] can be modulated by the activities of ecto-ATPases, ecto-AK and ecto-NDPK.

**Figure 4.**
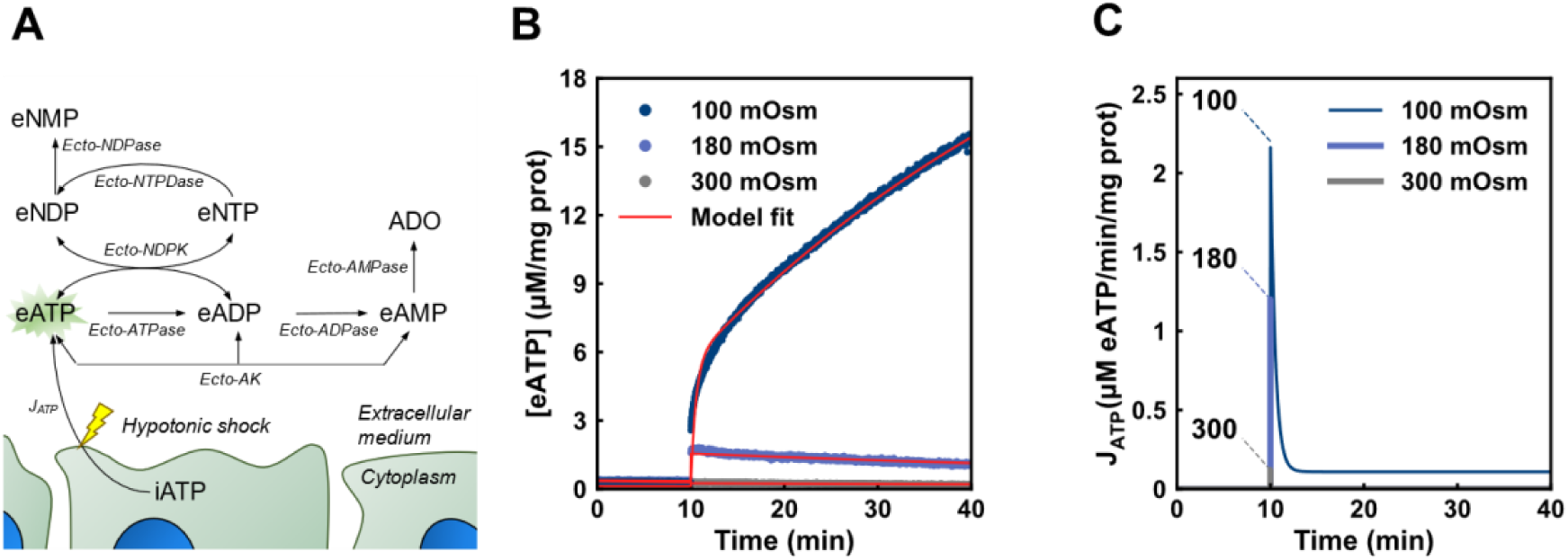
A model of extracellular purinergic regulation in non-polarized Caco-2 cells. (A) The scheme shows a representation of the model created to explain the experimental results in non-polarized cells. The yellow bolt indicates that the J_ATP_ depended on the application of a hypotonic shock. ADO means extracellular adenosine. The green star behind “eATP” indicates that this is the metabolite measured directly during experiments. (B) The plot shows, in red, the model fitting to eATP kinetics exposed to media of different osmolarities (experimental data correspond to those shown in Fig 1A and C). (C) iATP efflux (J_ATP_) predicted by the model upon the hypotonic or isotonic shocks indicated in the figure.

The model provides functions describing each of the fluxes involved in transport and metabolism of extracellular nucleotides (see S1 Table and section 4.13.1). Fitting the model to the experimental eATP kinetics under the different conditions allowed to obtain the best-fit values for the parameters of these functions (S1 Table). In that way the contributions of each flux to eATP kinetics were quantified, and several predictions were made.

##### 2.1.4.1. iATP release

For experiments under iso- and hypotonic media, the model found a good fit to experimental data (continuous lines in Fig 4B), thus allowing to predict the rate of iATP efflux (J_ATP_) over time (Fig 4C). J_ATP_ was rapid and transient in nature, leading to a 12-fold increase of [eATP] to a maximum in less than 2 seconds under the 180 mOsm shock, followed by rapid inactivation. The magnitude of the J_ATP_ peak depended on the osmotic gradient imposed. Inactivation of J_ATP_ was observed under conditions where no lysis was detected (isotonic and 180 mOsm media). On the other hand, a lytic flux (J_L_) explains the continuous increase of [eATP] at 100 mOsm (Fig 1C and 4B).

##### 2.1.4.2. Ecto-enzymes

Another factor shaping eATP kinetics is eATP hydrolysis by ecto-ATPase activity. We have previously observed that, in intact non-polarized Caco-2 cells, ecto-ATPase activity follows a linear function of micromolar concentrations of eATP [14]. Thus, following a stimulus promoting iATP release, any increase of [eATP] should be at least partially counterbalanced by an increase of ecto-ATPase activity.

Model predictions made at 180 mOsm show that the initial peak of [eATP] increase due to J_ATP_ is about 8-fold higher than the rate of eATP hydrolysis, *i.e.,* J_ATP_ was 1.2 µM iATP/min/mg of protein (Fig 4C) and eATP hydrolysis was 0.15 µM eATP/min/mg of protein at 1.5 µM eATP (S1 Table). Thus, during the first seconds of [eATP] increase, eATP kinetics was mainly governed by iATP release. At later times, however, the J_ATP_ inactivated, and the ecto-ATPase activity progressively gained importance in controlling [eATP]. This is illustrated by modelling a change in the amount of ecto-ATPase over a wide range, showing that a 5-fold increase of ecto-ATPase activity could lead to rapid decay of [eATP], while a 5-fold decrease would prolong high levels of [eATP] over the entire incubation period (Fig 5A). However, a similar procedure, *i.e*, increasing or decreasing 5 times the activity of ecto-AK, had no influence on the [eATP] during the hypotonic shock (not shown). This can be attributed to the sigmoidal kinetics of ecto-AK, whose activity is very low below 3 μM [eADP], but significantly higher above that concentration (Fig 5D). Thus, ecto-AK might influence [eATP] kinetics only when [eADP] is sufficiently high. Figure 5B shows a simulation where the initial [eADP] was raised up to 3 μM. At 3 μM [eADP], eATP degradation was comparable to eATP synthesis by ecto-AK, indicating that ecto-AK can counterbalance ecto-ATPase activity.

**Figure 5.**
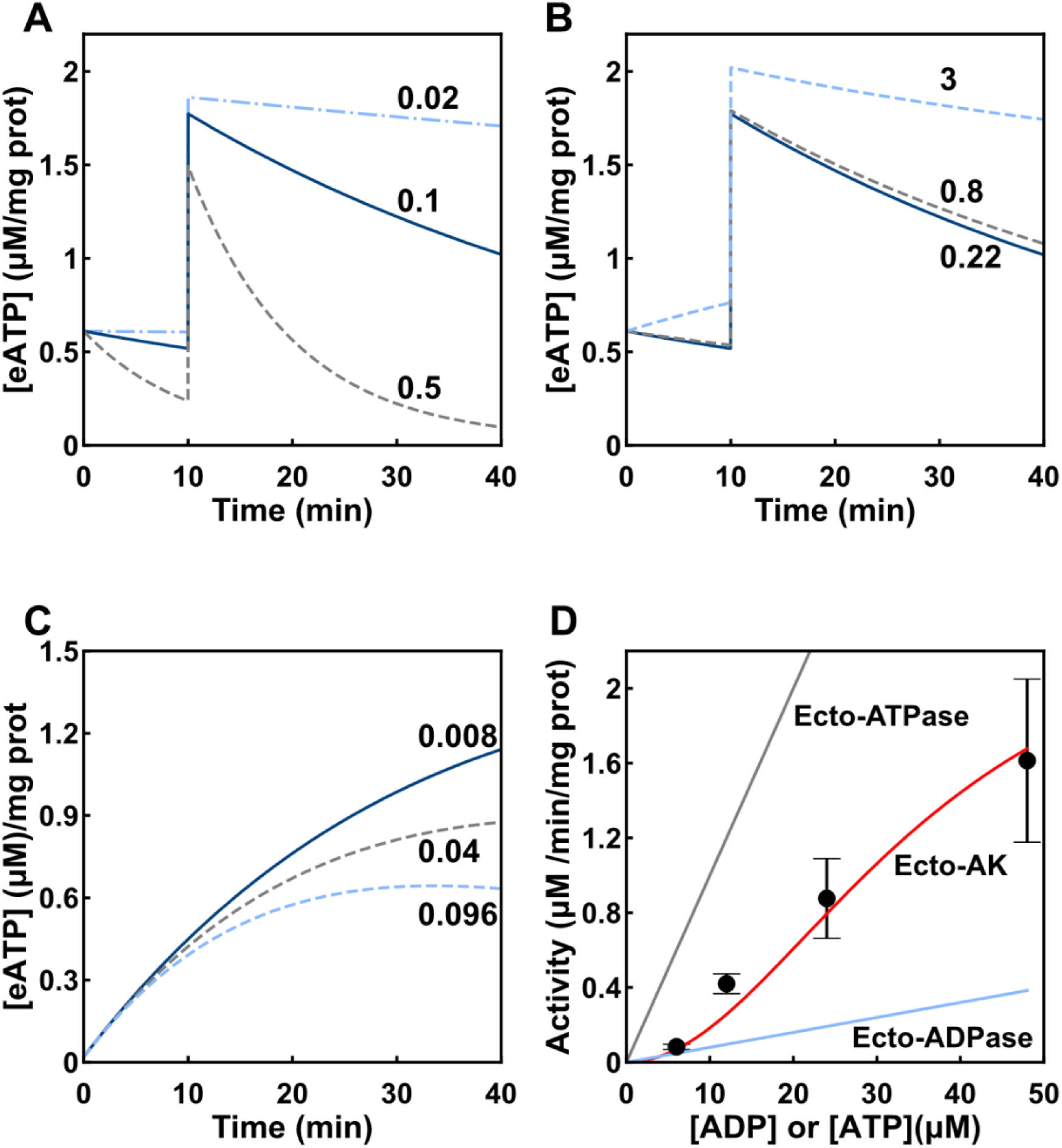
Role of ecto-AK, ecto-ATPase and ecto-ADPase activity on eATP dynamics. (A) The simulation shows the [eATP] as a function of time upon a 180 hypotonic shock when the ecto-ATPase activity displayed its measured value (0.1, continuous line in blue), a 5-fold increase (0.5, dashed line in grey), and a 5-fold decrease (0.02, dashed line in light blue). The numbers in the plot indicate the kinetic constant of the activity in (*μM eATP hydrolized*)/(*mg prot* / *μM eATP* / *min*) units. (B) The simulation shows the [eATP] as a function of time upon a 180 mOsm shock at different initial [eADP] concentrations: calculated pre-stimulus eADP (0.22 µM continuous line in blue), a 3.5-fold increase (0.8 µM, dashed line in grey), and a 14-fold increase (3 µM, dashed line in light blue). (C) The simulation shows the [eATP] as a function of time upon addition of 6 µM [eADP] (the corresponding experimental results are shown in Fig. 2A). The plot shows the eATP kinetics under various values of the kinetic constant for ecto-ADPase, i.e, the constant experimentally determined (0.008, continuous line in blue), a 5-fold increase (0.04, dashed line in grey), and a 12-fold increase (0.96, dashed line in light blue). The numbers in the plot indicate the kinetic constant of the activity in (*μM eADP hydrolized*)/(*mg prot* / *μM eADP* / *min*) units. (D) Ecto-ATPase, ecto-AK and ecto-ADPase activites as a function of their respective substrates, eATP for ecto-ATPase and eADP for ecto-AK and ecto-ADPase. The points show the initial velocities for eATP synthesis as a function of [eADP] by ecto-AK calculated from experimental data shown in Fig 2A. The points are means ± s.e.m. from 3 to 5 independent experiments run in duplicate. The continuous lines represent enzyme activities as a function of their respective substrates (see S1 Table for further details).

Another factor to consider is ecto-ADPase activity. We have previously shown that Caco-2 cells displays high ecto-ATPase but very low ecto-ADPase activity [14]. Nevertheless, a hypothetical increase of ecto-ADPase activity could negatively modulate ecto-AK activity. For example, an increase of 5- and 12-fold of ecto-ADPase activity would result in a 17% and 33% decrease in the [eATP] production respectively, at 6 µM [eADP] (Fig 5C).

Model predictions showed above implied that the expression of ecto-AK in Caco-2 cells may have an important role in [eATP] kinetics. To assess this hypothesis, we compared ecto-ATPase, ecto-ADPase and ecto-AK activities as a function of their respective substrate’s concentrations, that is, [eATP] for ecto-ATPase and [eADP] for ecto-ADPase and ecto-AK (Fig 5D). In Fig. 5D, symbols of ecto-AK activities represent the initial velocities for eATP synthesis as a function of [eADP] calculated from experimental data shown in Fig 2A, and the continuous line represents the fit to data of the ecto-AK function included in the model (details in S1 Table and in the work of Sheng and collaborators [17]). The ecto-ATPase and ecto-ADPase activities are predictions made from data of our previous work [14]. Ecto-ATPase displayed the highest rate of the three reactions. On the other hand, although at low [eADP], ecto-AK and ecto-ADPase activities are similar and have relatively low values, the sigmoidal kinetics of ecto-AK allows a strong activity increase as [eADP] is raised, thus reaching activity levels well above those of ecto-ADPase activity (Fig 5D).

Finally, in the presence of non-adenosine nucleotides, the influence of ecto-NDPK on eATP dynamics was assessed and analysed. Caco-2 cells synthetised eATP by ecto-NDPK activity in the presence of eCTP, eUTP and eGTP as NTP donors, and exogenous and endogenous eADP (Fig 3).

The model found a good fit to the experimental [eATP] kinetics in the presence of 100 μM eCTP and different concentrations of eADP (Fig 6A). Model predictions of ecto-NDPK activity at different [eADP] agreed well with initial velocities of experimental ecto-NDPK activities shown in Fig. 3A (Fig 6B). We also studied the effect of eUTP addition without the addition of exogenous eADP (a condition where only endogenous eADP was present, S3 Fig) on the transient rise of [eATP] (Fig 3C, replicated in Fig 6C). To understand the role of eNTPs on ecto-NDPK activity, it is important to recall that ecto-NTPDases of Caco-2 cells can hydrolyse non-adenine nucleotides (S4 Fig). Model predictions show changes in ecto-NDPK and ecto-ATPase activities (Fig 6D), and the corresponding dynamics of [eATP] and [eADP] (Fig 6E), and of [eUTP] and [eUDP] (Fig 6F). Kinetics of eATP (Fig 6C and E) could be analysed in 3 stages. First, [eATP] increases due to a high an ecto-NDPK/ecto-ATPase activities ratio in the presence of high [eUTP] and basal eADP (stage 1 in Fig 6D, E and F). The resulting elevated [eATP] activates ecto-ATPase activity, while ecto-NDPK decreases deeply because its substrates (eUTP and eADP) are consumed by ecto-NTPase activity and by ecto-NDPK activity itself. A balance is then established between ecto-NDPK and ecto-ATPase activities in stage 2, where [eATP] is transiently stable. Finally in stage 3, [eUTP] continues decreasing, leading to a high ecto-ATPase/ecto-NDPK activities ratio, causing [eATP] to decrease and [eADP] to rise again (Fig 6E).

**Figure 6.**
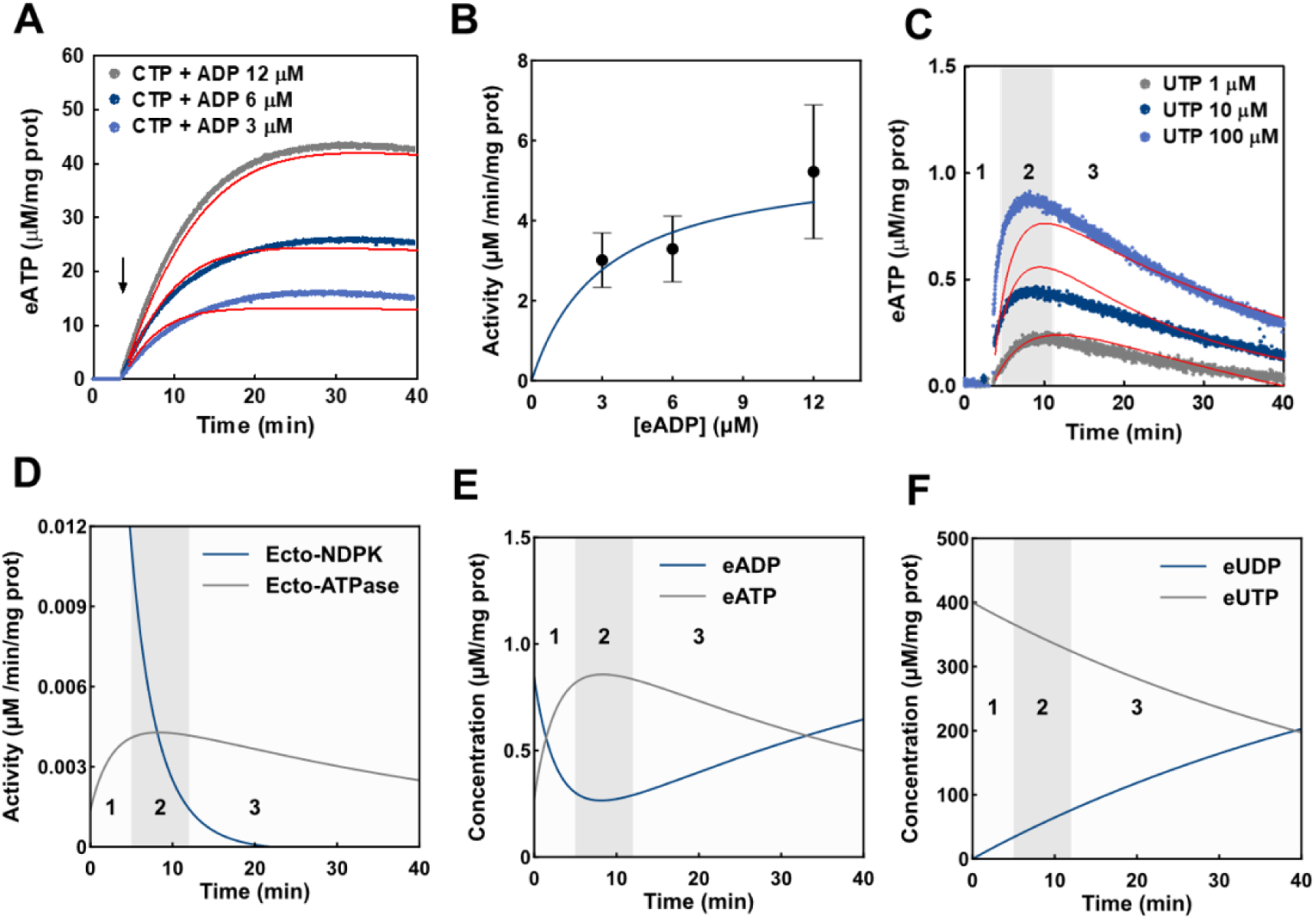
Role of ecto-NDPK on eATP dynamics. (A) The plot shows the experimental results of [eATP] dynamics in the presence of 100 µM [eCTP] and various [eADP] (also shown in Fig. 3 A). Model fitting was applied to data and shown as continuous red lines. (B) The plot shows the ecto-NDPK activity expressed in µM of ATP synthetised per minute per mg of protein. The dots represent the initial velocity of ecto-NDPK (calculated from the experimental data shown in panel A) as a function of [eADP]. Points represent the means ± s.e.m. from 3 independent experiments run in duplicate. The continuous line represents the ecto-NDPK activity predicted by the model (details in S1 Table). (C) The plot shows the experimental [eATP] dynamics in the presence of various [eUTP] concentrations (also shown in Fig 3C) and the continuous lines represent the model fitting. (D) The plot shows time changes of ecto-NDPK (blue line) and ecto-ATPase (grey line) activities predicted by the model. In the plot, the zones 1 (white background), 2 (grey background) and 3 (white background) represents the [eATP], increase, stabilization and decrease stages respectively. In (E) and (F) the plot shows the model predictions of [eATP] and [eADP], or [eUTP] and [eUDP] respectively as a function of time upon addition of 100 µM eUTP to non-polarized Caco-2 cells. Data is expressed in µM/mg protein, which was calculated by dividing the [eATP] at any time by the average protein mass in the experiments (0.25 mg in average).

### 2.2. eATP regulation in polarized Caco-2 cells

Because several reports showed differential activities of enzymes and transporters at each side of polarized epithelia [18, 19], we speculated that eATP regulation might be different at the apical and basolateral sides of polarized Caco-2 monolayers.

We then used polarized Caco-2 cells to test the effect of hypotonic shock on iATP release and resulting eATP kinetics at the apical and basolateral sides. Similarly to the procedure employed for non-polarized cells, we fitted the model shown in Fig 4A to the experimental data to understand quantitatively the mechanisms involved in [eATP] regulation in differentiated monolayers of Caco-2 cells.

Experimental results show that, following a 180 mOsm hypotonic shock, [eATP] increased at both sides of the monolayers, with qualitatively different kinetics. While at both sides the initial rate of [eATP] increase was fast, apical eATP kinetics achieved a maximum at 1.5 minutes, followed by a rapid decay. This was not observed in the basolateral domain, where [eATP] continued increasing at a progressively lower rate, and a very slow [eATP] decay was observed only after 20 minutes (Fig 7A).

**Figure 7.**
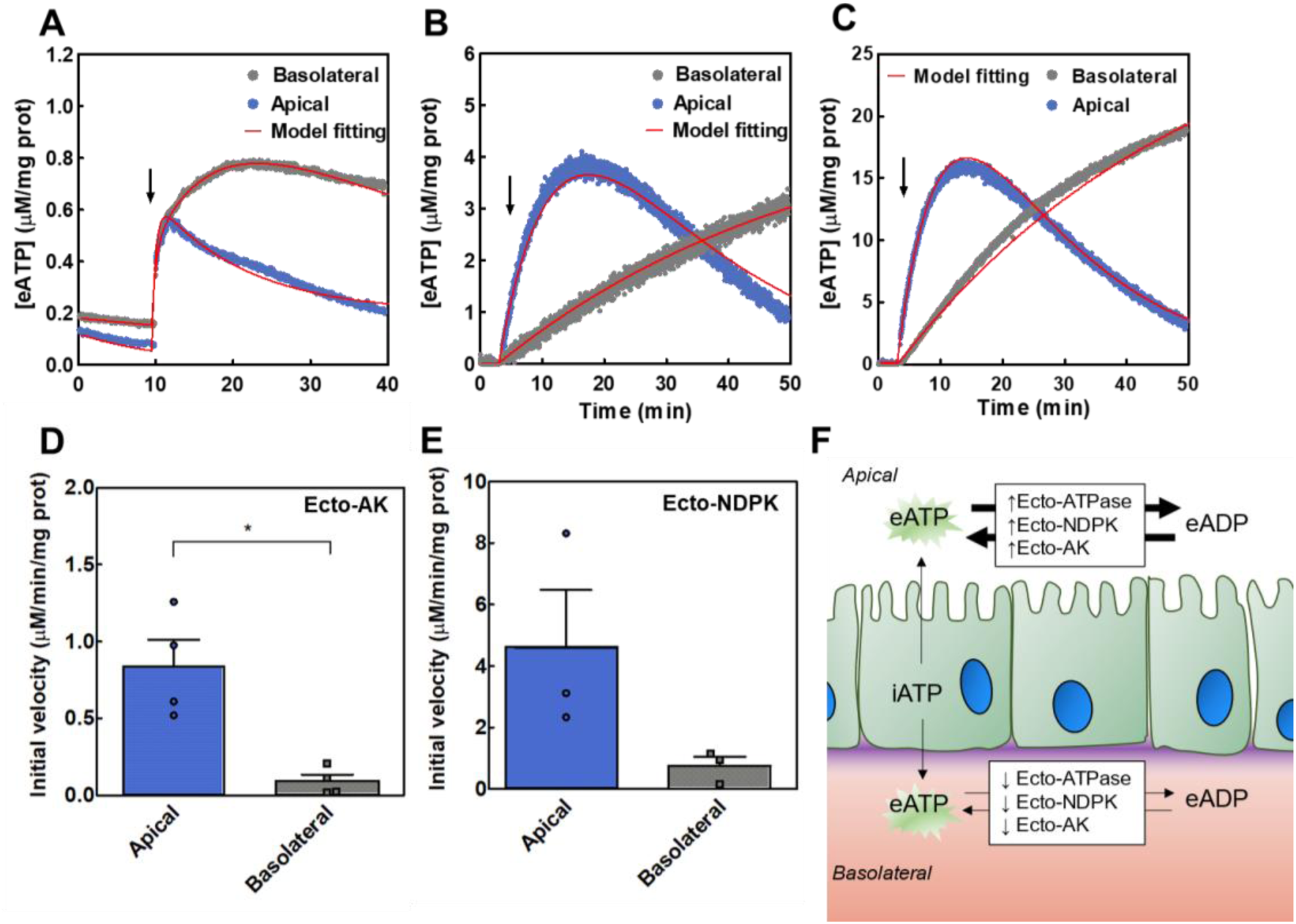
Apical and basolateral eATP regulation in Caco-2 monolayers. Results in A-C showed eATP kinetics in the apical and basolateral sides of polarized Caco-2 monolayers. Quantification of [eATP] was performed separately on each side of the monolayer. (A) Effect of hypotonic shock on eATP kinetics. At the times indicated by the arrow, cells were exposed to 180 mOsm medium on the basolateral (grey) or the apical (blue) compartments. Data are the means from 5 independent experiments. (B) Ecto-AK activity. eATP kinetics in the presence of 12 µM eADP added to the basolateral (grey) or apical (blue) compartments. Data are the means from 2 independent experiments. (C) Ecto-NDPK activity. eATP kinetics in the presence of 100 µM eCTP + 12 µM eADP added to the basolateral (grey) or the apical (blue) compartments. Experiments were run in the presence of 10 μM Ap5A (adenylate kinase blocker). Data are the means from 3 independent experiments. (D) Ecto-AK initial velocities in polarized Caco-2 cells. Data are means + s.e.m. of 4 independent experiments. * indicates a P-value < 0.05 in comparison with the apical condition. (E) Ecto-NDPK initial velocities in polarized Caco-2 cells. Data are means ± s.e.m. of 3 independent experiments. (F) Scheme of the results interpretation showing that the increased activity of Ecto-AK, Ecto-NDPK and Ecto-ATPase leads to a faster eATP turnover.

The two different eATP kinetics suggested different activities of ecto-enzymes present at both sides of the monolayers. Therefore, we determined the activities of ecto-ATPase, ecto-AK and ecto-NDPK.

For assessing ecto-ATPase activity, polarized Caco-2 cells were exposed to various [eATP] (0.2 – 7 µM) and eATP hydrolysis was estimated by quantifying [eATP] decay rates (S5 Fig). The initial rate values of [eATP] decay were used to calculate ecto-ATPase activity at each [eATP], so as to build a substrate curve (S5C Fig). Linear fitting to experimental data showed that ecto-ATPase activity was 4-fold higher in the apical than in the basolateral domain.

To assess ecto-AK activity, Caco-2 cells were exposed to 12 µM eADP at the basolateral or apical domains. In the apical domain, [eATP] increased rapidly to a maximum, followed by a rapid decay towards pre-stimulated levels, while basolateral [eATP] increased steadily at a lower rate (Fig 7B). Initial velocity estimations showed that ecto-AK activity was significantly higher in the apical than in the basolateral compartment (Fig 7D).

Ecto-NDPK activity was quantified using polarized cells exposed to 100 µM eCTP plus 12 µM eADP in the basal and apical domains. Experiments were run in the presence of 10 μM Ap5A to block ecto-AK activity. Production of [eATP] by ecto-NDPK was much higher than that observed under conditions used to measure ecto-AK activity, though the domain specific pattern of eATP kinetics was similar when assessing the two ecto-kinases, *i.e*., a biphasic pattern in the apical domain, and a steady [eATP] increase, at a lower rate, in the basal domain (Fig 7C). The initial velocity of ecto-NDPK was higher in the apical than in the basolateral domain although differences were not significant (*p* value = 0.1).

A good fitting of the model to all experimental data was achieved (red lines in Figs 7 A-C). The model fitting allowed to obtain the ecto-NDPK and ecto-AK maximal velocity (Vmax) and compared them with the ones obtained from non-polarized cells (S6 Fig and S2 Table). Results indicated that the ecto-NDPK maximal activity in the apical compartment is a slightly higher than that of the non-polarized cells and significantly higher than that of the basolateral compartment. On the other hand, the ecto-AK maximal activity is significantly higher compared with the basolateral compartment or the non-polarized cells. Thus, the differences between the apical and basolateral eATP dynamics can be explained by an increase or decrease in ecto-enzymes activities.

Altogether experimental results showed significantly higher activities of the ecto-enzymes (ecto-ATPase, ecto-AK and ecto-NPDK) in the apical, as compared to the basolateral domain (Fig 7F).

### 2.3. Ecto-AK and ecto-NDPK are active in human primary small intestinal epithelial cells

Having characterized ecto-AK and ecto-NDPK activities of Caco-2 cells, we wondered whether these ecto-enzymes would be functional in IECs extracted from human small intestine. Accordingly, we used samples obtained from small intestine biopsies from healthy donors (Fig 8).

**Figure 8.**
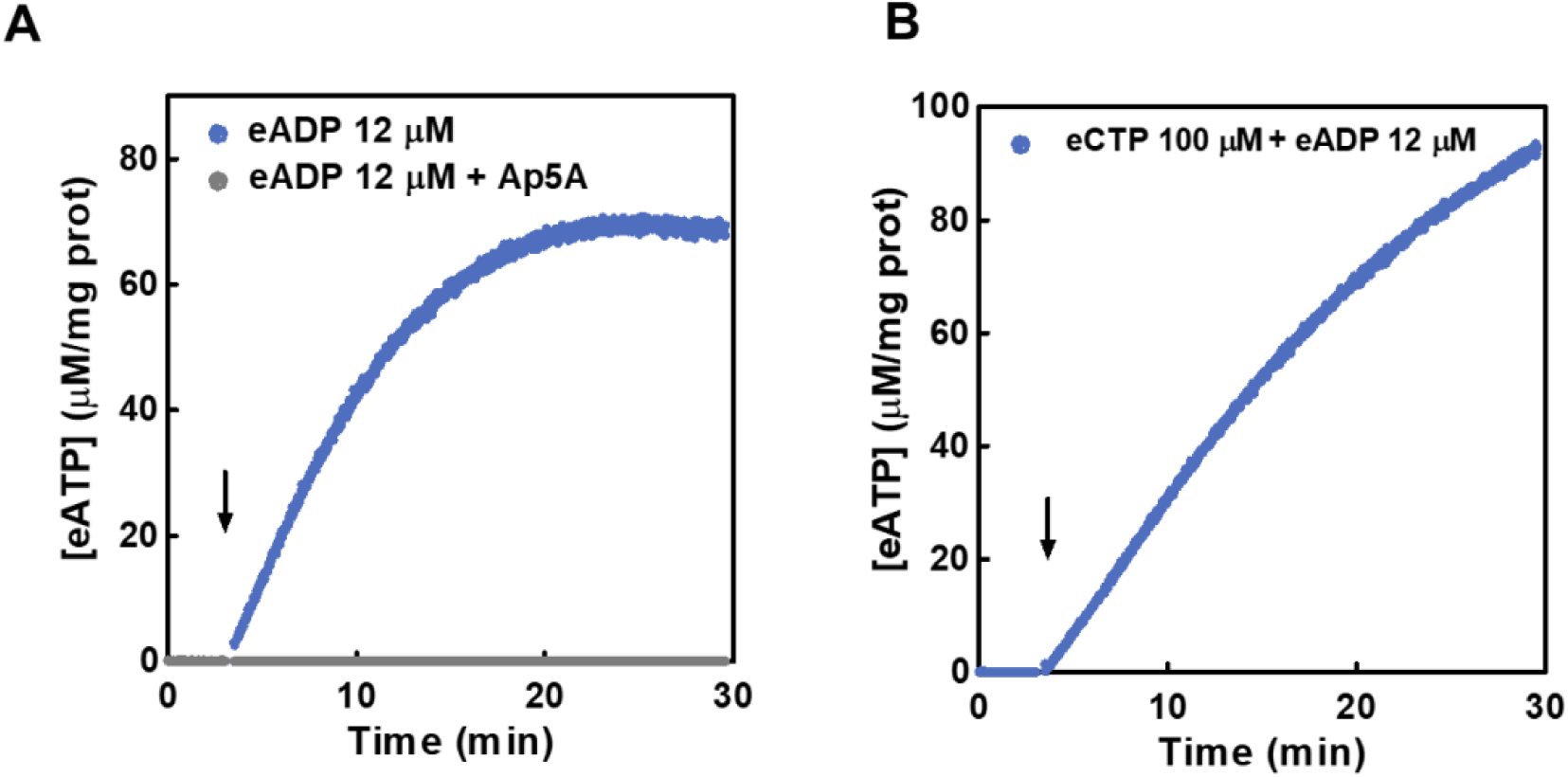
Ecto-AK and ecto-NDPK activities in IECs. Time course of eATP synthetized from exogenous eADP (A) or eCTP and eADP (B) in the extracellular medium of IECs. (A) The cells were incubated with the luciferase-luciferin reaction mix and 12 μM eADP was added at the time indicated by the arrow in presence (grey) or absence (blue) of 10 μM Ap5A (B) 100 μM eCTP plus 12 μM eADP, in the presence of 10 μM Ap5A were added at the time indicated by the arrow. [eATP] was quantified by luminometry. Values are the means of 3 independent experiments run in duplicate.

Results show that exposure of IECs to both 12 µM eADP (to assess ecto-AK) (Fig 8A) or to 12 µM eADP + 100 µM eCTP (to assess ecto-NDPK, Fig 8B) led to significant eATP production in the micromolar range. Furthermore, as expected, the presence of ApA5 totally inhibited the ecto-AK activity (Fig 8A).

## 3. Discussion

Intestinal epithelial cells can release iATP and express several ecto-enzymes capable of regulating the amount and metabolism of eATP at the cell surface. The main goal of this study was to characterize quantitatively the dynamic interplay of iATP release, eATP hydrolysis and eATP synthesis contributing to the dynamic regulation of [eATP] in Caco-2 cells. Special emphasis was given to the role of ecto-kinases promoting eATP production under different conditions.

Since Caco-2 cells undergo spontaneous enterocytic differentiation in culture, we decided to first approach the complexity of eATP regulation using the relatively simpler non-polarized cell model, and later extend the study to fully differentiated cells. These form apical and basolateral poles where morphological and biochemical features are segregated [18].

When exposed to hypotonicity, non-polarized Caco-2 cells triggered a strong iATP efflux that rapidly inactivated, leading to low µM eATP accumulation. A number of studies have confirmed that such micromolar [eATP] are capable of activating P2 receptors with high affinity for that nucleotide, such as P2Y2, P2Y11 and almost all P2X receptors [20]. In Caco-2 cells, eATP dose dependently activates P2Y receptors involved in the activation of MAPK cascades and transcription factors that promote cell proliferation [21, 22].

In principle, purinergic activation by eATP should be transient, due to the presence of ecto-nucleotidases, the activities of which promotes strong eATP hydrolysis in Caco-2 cells [14]. Accordingly, our results show that hypotonicity induced iATP release and concomitant [eATP] accumulation, where [eATP] decay was accelerated by constitutive ecto-ATPase activity. This decay was even higher for a model predicted upregulation of eATP hydrolysis by one or more ecto-nucleotidases, as occurs in various cells and tissues during pathogen infection [23], cell differentiation [24] or tumorigenesis [25].

The above results imply that iATP release and eATP hydrolysis constitute two opposing fluxes shaping eATP kinetics of Caco-2 cells. However, the presence of ecto-kinases found in this study suggest that the dynamic regulation of [eATP] should also take the activities of these enzymes into account.

In this respect, addition of exogenous eADP to Caco-2 cells dose dependently increased [eATP]. The fact that eATP synthesis was almost fully blunted by Ap5A, an AK blocker that does not permeate intact cells, suggested the presence of a functional ecto-AK. Results of the mathematical model allowed to envisage the contribution of ecto-AK to eATP kinetics. In the absence of exogenous eADP, the contribution of ecto-AK to eATP kinetics was negligible, so that [eATP] depended mainly on the balance between the rates of iATP release and eATP hydrolysis. This is due to the low endogenous [eADP] present under the experimental conditions. However, due to the sigmoidal nature of the AK reaction, model predictions show that increasing [eADP] in the low micromolar range, suffices to promote significant eATP synthesis by ecto-AK, upregulating eATP kinetics. Thus, under certain conditions, *e.g.,* when cell leak intracellular ADP (iADP) or eADP is supplied paracrinally by other cell types, eATP synthesis by ecto-AK of Caco-2 cells will transiently stabilize eATP levels, thereby favouring propagation of eATP-dependent purinergic signalling. A similar stabilizing role of ecto-AK on [eATP] has been proposed for HT29 cells, lung epithelial cells and lymphocytes [11,12,26].

Modelling shows that ecto-ADPase activity, which facilitates eADP degradation, may compete with ecto-AK for the available eADP. However, Caco-2 cells - as HT29 cells [12]-displayed a relative low ecto-ADPase activity, in agreement with the presence of a functional ecto-NTPDase 2 in both cell types [12, 14], and in addition the intrinsic sigmoidal nature of ecto-AK activity makes ecto-AK more sensitive to [eADP] than ecto-ADPase.

Another consequence of ecto-AK activation relates to P1 signalling, since activity of this enzyme will provide eAMP from eADP for further hydrolysis to adenosine by ecto-5’NT present in Caco-2 cells [14], finally leading to extracellular adenosine accumulation.

Our model predictions show how increasing [eADP] in the low µM range might lead to substantial adenosine accumulation, which may engage 4 different P1 receptors [27]. The consequences of P1 signalling on proliferation of Caco-2 cells and several other intestinal epithelial cell lines have been studied before [28]. In general, the balance between P1 and P2 receptors on epithelial cells regulate intestinal secretion [29–32] and absorption [33, 34]; responses triggered by the P2 receptor stimulation by eATP and other nucleotides are sometimes counteracted by P1 receptor stimulation by adenosine, though the potential role of ecto-AK was not considered in this context.

Another factor affecting eATP kinetics is ecto-NDPK. Activity of this enzyme was detected in many cells and tissues such as astrocytoma cells [35], endothelial cells [36, 37], lymphocytes [36], keratinocytes [38] and hepatocytes [39]. In general, ecto-NDPK will primarily serve to transfer phosphate groups between different extracellular nucleotides and thus potentially alter the pattern of P2 receptor activation. This is especially important since P2 receptor subtypes are differentially selective for adenine and uridine eNDPs and eNTPs [40, 41].

Our results show that ecto-NDPK can use eCTP, eGTP and eUTP to phosphorylate eADP to eATP. As model predictions show, activities of ecto-NDPK (promoting eATP synthesis from eUTP and eADP) and ecto-nucleotidase (promoting eATP and eUTP hydrolysis) change in opposite directions to transiently stabilize [eATP].

Results analysed above show that, in non-polarized Caco-2 cells, [eATP] can increase by iATP release and ecto-kinase mediated eATP synthesis and decrease by ecto-nucleotidases mediated by eATP hydrolysis.

Next, we studied [eATP] dynamics of polarized Caco-2 cells. These cells differentiate spontaneously into polarized cells, with apical and basolateral domains exhibiting morphological and biochemical features of small intestine enterocytes [18, 42]. In particular, the Caco-2 polarized phenotype is characterized by high levels of hydrolases typically associated with the brush border membrane. The fact that in a variety of epithelia several ecto-nucleotidases and ecto-phosphatases preferentially - but not exclusively-locate in the apical domain [43–45], anticipated a different eATP regulation at both poles of Caco-2 cells.

Accordingly, hypotonically induced eATP kinetics had a faster resolution and was more effectively regulated at the apical, than at the basolateral side, a result in agreement with the observed higher apical (than basolateral) ecto-ATPase activity measured in this study. This is in agreement with several reports using intestinal epithelial cell from murine models and human intestinal cell lines, showing that various isoforms of ecto-NTPDases, ecto-phosphatases and ecto-NPPases are preferentially located in the apical domain [45].

A qualitatively similar pattern was observed for ecto-AK and ecto-NDPK of Caco-2 cells, in that apical activities were much higher. Interestingly, the model describing eATP dynamics of non-polarized cells could be successfully fitted to eATP kinetics on each of the polarized domains, thus allowing to calculate the consequences of ectoenzymes sorting on eATP regulation.

The fact that the apical domain exhibited a higher turnover of extracellular nucleotides, leading to higher eATP regulation may have adaptive value, considering that iATP release is a common response of epithelial intestinal cells to enteric pathogens [46]. Extracellular ATP may then act as a danger signal controlling a variety of purinergic responses aimed at defending the organism from a variety of pathogens and their toxins present in the intestinal lumen.

## 4. Materials and methods

### 4.1. Chemicals

All reagents were of analytical grade. Bovine serum albumin (BSA), malachite green, adenosine 5′-triphosphate (ATP), adenosine 5′-diphosphate (ADP), cytidine 5′-triphosphate disodium salt (CTP), adenosine 5′-monophosphate (AMP), uridine 5’-triphosphate (UTP), uridine 5’-diphosphate (UDP), guanosine-5’-triphosphate (GTP), phosphate-buffered saline (DPBS), 4-(2-hydroxyethyl)-1-piperazineetahnesulfonic acid (HEPES), ammonium molybdate, Triton X-100, phenylmethylsulphonyl fluoride (PMSF), pyruvate kinase, phosphoenol-pyruvate (PEP), luciferase, coenzyme A and P1,P5-Di (adenosine-5′) pentaphosphate pentasodium salt (Ap5A) were purchased from Sigma-Aldrich (St Louis, MO, USA). D-luciferin was purchased from Molecular Probes Inc. (Eugene, OR, USA).

### 4.2. Solutions

In the experiments to measure eATP by luminometry (section 4.5), cells were incubated with isotonic buffer called isosmotic DPBS (300 mOsm) containing: 137 mM NaCl, 2.7 mM KCl, 1 mM CaCl_2_, 2 mM MgCl_2_, 1.5 mM KH_2_PO_4_ and 8 mM Na_2_HPO_4_, pH 7.4 at 37°C (assay medium). When applying a hypotonic shock to cells, the medium was changed for other containing the same components but with a lower NaCl concentration. Thus, DPBS with 100, 150 and 180 mOsm were prepared. The osmolarity of all media was measured with a vapor pressure osmometer (5100B, Lugan, USA)

When measuring phosphate (section 4.8.4) the following medium without phosphate was employed instead of isotonic buffer: 145 mM NaCl, 5 mM KCl, 1 mM CaCl_2_, 10 mM HEPES, and 1 mM MgCl_2_, pH 7.4 at 37°C.

### 4.3. Caco-2 cell culture

Caco-2 cells (ATCC, Molsheim, France) were grown in Dulbecco’s modified Eagle’s medium (DMEM-F12, Gibco, Grad Island, NY, USA) containing 4.5 g/L glucose (Sigma-Aldrich, St Louis, MO, USA) supplemented with 10% v/v fetal bovine serum (Natocor, Córdoba, Argentina), 2 mM L-glutamine (Sigma-Aldrich, St Louis, MO, USA), 100 U/mL penicillin, 100 μg/mL streptomycin and 0.25 μg/mL fungizone (Invitrogen, Carlsbad, CA, USA) in a humidified atmosphere of 5% CO_2_ at 37°C. For eATP kinetics measurements cells were directly seeded on glass coverslips. For ecto-nucleotidase activity experiments using the malachite green method, cells were seeded in cell culture 24-well plates (Corning Costar, NY, USA)

#### 4.3.1. Polarisation of Caco-2 cells

For preparation of polarized Caco-2 monolayers, cells were seeded in permeable supports (inserts) made of polyester (Transwell; 0.1 µm pore size, 1.12 cm^2^ cell growth area; Jet Biofil, China) in 12-well plates at a density of 3 × 10^4^ cells/0.5 mL per insert. The medium was changed after 3 days, and then after every 3 or 4 days. The polarized Caco-2 monolayers were used for experiments after the transepithelial electrical resistance reached a plateau (approximately 21 days after seeding). In polarized and non-polarized cultures contamination (including Mycoplasma) was routinely tested.

### 4.4. Human Intestinal Epithelial Cells (IECs) isolation

IECs were isolated from ileum biopsies collected from healthy volunteers who were endoscopically evaluated for colon cancer (N = 3) at the Favaloro Foundation University Hospital. Samples of non-tumoral, non-injured intestinal biopsies were collected and transported in ice-cold Hanks’s balanced salt solution (HBSS) for immediate processing. The biopsies were incubated in 5 mM ethylenediaminetetra-acetic acid (EDTA) and 1.5 mM dithiothreitol HBSS with agitation for 25–30 minutes at room temperature to obtain IECs. Cells were pelleted, re-suspended in DPBS and used immediately.

The protocol for handling samples was approved by the Institutional Review Board of the Favaloro Foundation University Hospital (DDI [1587] 0621) and has been performed in accordance with the ethical standards laid down in the declarations of Helsinki and Istanbul. Informed consent was obtained from donors.

### 4.5. ATP measurements

The eATP concentration ([eATP]) of non-polarized Caco-2, polarized Caco-2 monolayers or IECs was measured using the firefly luciferase reaction (EC 1.13.12.7, Sigma-Aldrich, St Louis, MO, USA), which catalyses the oxidation of D-luciferin in the presence of ATP to produce light [47]. As described below, using this method it was possible to determine eATP kinetics, the iATP content and the activities of ecto-enzymes. Before the experiments, the cells were washed two times with the assay medium (isosmotic DPBS with or without Pi).

In this work, the cells’ medium was substituted by the assay medium before any measurement, therefore exoenzymes (enzymes release to extracellular medium not bound to the membrane) were removed and only ecto-enzymes (membrane bound extracellular enzymes) were investigated.

### 4.6. eATP kinetics of non-polarized Caco-2 and IECs

Non-polarized Caco-2 cells and IECs were seeded on glass coverslips. Under all conditions cells were mounted in the assay chamber of a custom-built luminometer, as previously described [48]. Because luciferase activity at 37°C is only 10% of that observed at 20°C [49], to maintain full luciferase activity, [eATP] measurements were performed at room temperature. The setup allowed continuous measurements of [eATP] by the luciferin-luciferase reaction.

A calibration curve was used to transform the time course of light emission into [eATP] versus time. Increasing concentrations of eATP from 13 to 1000 nM were sequentially added to the assay medium from a stock solution of pure ATP dissolved in isosmotic or hypotonic medium, according to the experiment. Calibration curves displayed a linear relationship within the range tested. After each experiment, cells were lysed with a solution containing 1 mM PMSF and 0.1% of Triton X-100 and the protein contents of each sample were quantified [50]. Results were expressed as [eATP] at every time point of a kinetics curve denoted as “eATP kinetics”, with [eATP] expressed as μM of eATP/mg protein in a final assay volume of 100 μL.

### 4.7. eATP kinetics of polarized monolayers

Polarized Caco-2 cells monolayers were placed in the insert physically separating an apical and a basolateral compartment. Detection of eATP was performed separately on either side, by adding the luciferin-luciferase mixture in one compartment (apical or basolateral) and adding isosmotic DPBS to the other side. In preliminary experiments, we observed that the luciferin-luciferase mix added in one compartment did not cross the monolayer into the other compartment. Thus, luminescence registered when measuring the [eATP] in one compartment was not contaminated by light from the other compartment due to luciferin-luciferase leakage.

When an hypoosmotic shock was applied, a luciferin-luciferase mix in DPBS with an osmolarity of 180 mOsm was added to the compartment of interest while, isosmotic DPBS was added to the other side.

### 4.8. Activities of ecto-enzymes

Ecto-ATPase, ecto-AK and ecto-NDPK activities of intact cells were measured by luminometry (section 4.5). Ecto-nucleotidase activities were measured by measuring the inorganic phosphate (Pi) release.

#### 4.8.1. Ecto-ATPase activity

Cells were exposed to different [eATP] (0.2, 1.2, 4.2 or 7 μM). Following an acute increase of [eATP], ecto-ATPase activity was estimated from the initial velocity of eATP decay at each [eATP].

#### 4.8.2. Ecto-AK activity

Cells were exposed to different [eADP] (6, 12, 24 or 48 μM) and the eATP kinetics was quantified in the absence and presence of 10 µM Ap5A (an AK blocker). Ecto-AK initial velocity was estimated as indicated in section 4.12.

#### 4.8.3. Ecto-NDPK activity

Cells were exposed to different [eADP] (3, 6 or 12 μM) in the presence of eCTP (100 μM), eGTP (100 μM) or eUTP (1, 10 or 100 μM). Then, the eATP kinetics was quantified in the presence of Ap5A to block the eADP to eATP conversion by ecto-AK activity. In some experiments 5 mM eUDP was added to inhibit ecto-NDPK activity. Ecto-NDPK initial velocity was estimated as indicated in section 4.12.

#### 4.8.4. Ecto-NTPDase activities

Cells were incubated with 500 μM of eCTP, eUTP or eGTP at 37°C. Samples were taken at 30, 60, 90 and 120 minutes after nucleotides addition and, the inorganic phosphate concentration was measured by the malachite green method [14, 51].

Activities measured in section 4.8.1 were expressed as µM of eATP hydrolysed per minute, normalized by the cell protein mass in the experimental sample (µM of eATP /mg protein/min). Results from experiments explained in sections 4.8.2 and 4.8.3 were expressed as µM of eATP synthetised per minute, normalized by the cell protein mass in the experimental sample (µM of eATP /mg protein/min). Activities measured in section 4.8.4 were expressed as µM of inorganic phosphate released per minute, normalized by the cell protein mass in the experimental sample (µM of Pi /mg protein/min)

### 4.9. Intracellular ATP measurements

Caco-2 (0-30,000 cells) were laid on coverslips, incubated with 45 μL of luciferin-luciferase reaction mix for 5 minutes and subsequently permeabilized with digitonin (1.6 mg/mL final concentration). Light emission was transformed into eATP concentration as a function of time as indicated in section 4.6. After considering the total volume occupied by Caco-2 present in the chamber, and the relative solvent cell volume (3.66 µl per mg of protein) [52], [iATP] was calculated in mM. To calculate the % of iATP release, the following equation was employed:

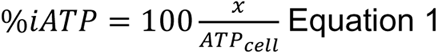

where ATP_cell_ represents the [ATP] obtained when iATP from all cells is released into the assay medium. The “x” denotes the [eATP] measured at any time. The value of ATP_cell_ was 66 µM/mg protein and was calculated by multiplying the [iATP] (1.8 mM, section 2.1.1) by the Caco-2 cell volume (3.66 µl per mg of protein [52]) and diving by the assay volume (0.1 mL).

### 4.10. Extracellular ADP measurements

For detection of extracellular ADP (eADP) of intact Caco-2 cells, 3 U/100 µl of pyruvate kinase and 100 μM PEP were added to the luciferin-luciferase mix. Using PEP as a substrate, pyruvate kinase promotes the stoichiometric conversion of eADP into eATP [53]. The resulting eATP was then measured by light emission using the luciferin-luciferase procedure described above.

### 4.11. Data analysis

Statistical significance was determined using the non-parametric Mann-Whitney test. Data were analyzed and graphically represented using GraphPad Prism software v5.0 (Graph Pad Software, San Diego, CA, USA). Each independent experiment was carried out in an independent cell culture or tissue sample in a different day.

### 4.12. Initial velocity estimation

To measure the initial velocity of Ecto-AK or Ecto-NDPK, the eATP dynamics were measured as indicated in section 4.8.2 and 4.8.3. Only the values of [eATP] obtained during the first 5 minutes after substrates addition were considered for further analysis. The following equation was fitted to experimental data:

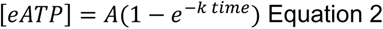

where A and k are parameters, whose value are optimized to achieve a good fitting of Eq. 2 to experimental data. The initial velocity is the derivative of [eATP] as a function of time at time 0 (the time when substrates were added). Thus, the initial velocity was calculated by multiplying the value of A by the value of k.

### 4.13. Mathematical modelling

Chemical models of extracellular nucleotides were built using COPASI (Complex Pathway Simulator) software in version 4.29 (source: https://copasi.org/) [54]. Parameter optimization was performed using COPASI “parameter estimation function” with Hooke & Jeeves, Levenberg-Marquardt, or Evolutionary programming as optimization methods. An initial guess of the parameter value was proposed based on literature data for each kinetic step. A detailed description of the models employed in this work can be found in S1 and S2 Tables. Parameters obtained from the model fitting are expressed as the best value ± standard deviation. The COPASI files of the models described in section 4.13.1 and 4.13.2 can be found in the data repository (see data availability statement).

#### 4.13.1. A model of purinergic homeostasis in non-polarized Caco-2 cells

To explain the experimental observations, a data driven mathematical model was created (depicted in Fig. 4A). The model has 7 reactions to explain the chemical fluxes of transformations or transport of extracellular nucleotides in Caco-2 cells: J_ATP_, J_Ecto-ATPase_, J_Ecto-ADPase_, J_Ecto-AMPase_, J_Ecto-AK_, J_Ecto-NDPK_ and J_Ecto-NTPDase_. A detailed description of each flux, its mathematical description and parameters can be found in S1 Table. In the model, the concentration of each species as a function of time was calculated from the following differential equations:

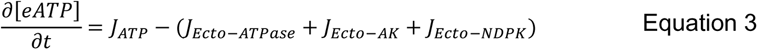

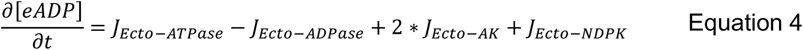

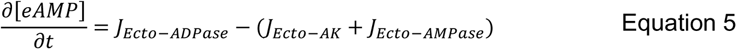

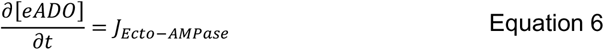

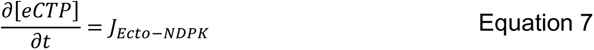

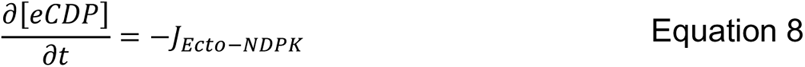

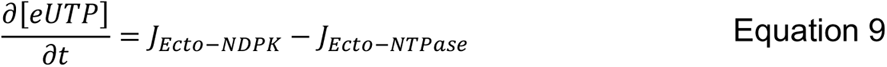

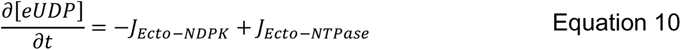

Note that in the equations J_Ecto-AK_ and J_Ecto-NDPK_ were considered in the direction of eATP consumption, *i.e.,* eAMP + eATP ↔ 2 eADP for J_Ecto-AK_ and eNDP + eATP ↔ eNTP + eADP for J_Ecto-NDPK_. The model was written in COPASI 4.29 and was fitted simultaneously to all experimental data shown in Fig. 1 A, Fig. 1 C, Fig. 2 A, Fig. 3 A and Fig. 3 B. The fitting of the model to experimental data can be seen in Figs. 4B, 5C, 6A and 6C as red lines.

Some kinetic parameters of the enzymes catalyzing the reactions were obtained from the literature. Parameters from the J_ecto-ATPase_ and J_ecto-ADPase_ were obtained from our previous work [14]. The V_max_ of the J_ecto-AMPase_ reaction was obtained from our previous work [14], while the K_m_ was obtained from the work of Navarro et al. [55]. Kinetic parameters of the J_ecto-AK_ activity were obtained from the work of Sheng *et al.* [17]. The equilibrium constant (K_eq_) and the affinity for ATP (K_mAT_) of the J_ecto-NDPK_ were obtained from the work of Garces and Cleland [56]. The affinity constants for product inhibition in J_ecto-NDPK_ (K_iNDP_ and K_iADP_) were estimated from the work from Lascu *et* Gonin [57]. The rest of the model parameters were obtained from model fitting to experimental data (see S1 and S2 Tables for more details). The shape of the J_ATP_ flux as a function of time was modeled based on findings of a previous work from our group [58].

#### 4.13.2. A model of purinergic homeostasis in polarized Caco-2 cells

The model fitted to experimental data from the apical and basolateral compartments data is the same model indicated in section 4.13.1, although the parameters of some reactions were fitted again (S2 Table). The J_ATP_ expression for the 180 mOsm hypotonic shock in the polarized cells was different from the one employed on non-polarized cells (S2 Table). The mathematical expressions of the other 6 reactions were not modified. Four parameters were refitted to the data to account for differences in the ecto-ADPase, ecto-AK and ecto-NDPK activities after polarization (values can be found in S2 Table). Moreover, in the case of ecto-NTPDase, the eCTP hydrolysis could not be neglected in the apical compartment and was necessary to achieve a good fit to experimental data. In contrast the eCTP hydrolysis could be avoided in the basolateral compartment without affecting model fitting. This suggest that the ecto-CTPase activity is greater in the apical than in the basolateral compartment, in agreement with the increased activity of other enzymes on the apical side.

The differential equations for [eCTP] and [eCDP] are modified in the apical side model to account for the eCTP hydrolysis:

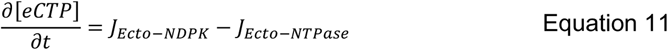

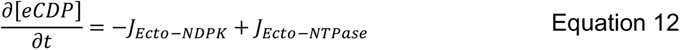

The models for the apical and basolateral compartments were written in COPASI 4.29 and fitted to experimental data shown in Fig. 7A, B and C. The COPASI files can be found in the data repository (see data availability statement).

## Abbreviation

eATP: Extracellular ATP
iATP: Intracellular ATP
eADP: Extracellular ADP
eAMP: Extracellular AMP
eUTP: Extracellular UTP
eUDP: Extracellular UDP
eCTP: Extracellular CTP
eCDP: Extracellular CDP
eGTP: Extracelullar GTP
P2 receptors: Purinergic receptor 2
Ecto-NTPDase: Ecto-nucleoside triphosphate diphosphohydrolase
Ecto-AK: ecto-adenylate kinase
Ecto-NDPK: ecto-nucleoside diphosphate kinase
NDP: nucleoside diphosphate
NTP: nucleoside triphosphate
mOsm: Mili osmol / litre
Pi: Inorganic phosphate
PEP: Phosphoenolpyruvate

Nucleotides expressed in brackets means concentration of that nucleotide, for example, [eATP] means extracellular ATP concentration.

When the word exogenous is employed, it means that the nucleotide was added from an external source and not synthetised or released by the cells. When the word endogenous is employed, it means that the extracellular nucleotide was release or synthesized by the cells.

## Acknowledgements

We are thankful to Dr. Cafferata for providing the Caco-2 cells.

## Data availability

Data can be found in the following doi:

10.6084/m9.figshare.21938651

or temporarily in the following link:

https://figshare.com/s/1fab1cb9fa543e8520c5

## Supporting information

**S1 Figure. iADP release estimation.** Increase in [eATP] after a 180 mOsm hypotonic shock in absence (blue) or presence (grey) of PK (3 U) and PEP (100 μM) were evaluated as ΔATP, i.e., the difference between [eATP] at 1 min post-stimulus and basal [eATP].

**S2 Figure**. **Inhibition by exogenous eUDP of ecto-NDPK activity in the presence of eCTP and eADP**. Time course of eATP accumulation in the presence of 100 eUTP μM in the absence (blue) or in the presence of 5 mM eUDP (grey). The data showed are the means of 3 independent experiments.

**S3 Figure. Measurement of eADP by the conversion to eATP.** Caco-2 cells were incubated with luciferin-luciferase and, at the time indicated with the arrow, PK (3 U) and PEP (100 μM) were added. The value of the [eADP] in resting conditions was 0.77 ± 0.44 µM eADP/mg. Given a usual protein cell mass of 0.2 mg, the [eADP] in resting conditions is 0.15 ± 0.09 µM. The data showed are the means of 5 independent experiments.

**S4 Figure. Ecto-nucleotidase activity of Caco-2 cells.** Experiments were performed in assay medium without Pi at room temperature, and Pi production was measured by the malachite green method (section 2.8.4). The time course of Pi accumulation in the extracellular media of Caco-2 cells was measured and values of enzyme activity were derived from initial rates of nucleotides hydrolysis for 500 μM of eUTP (grey), eGTP (blue) and eCTP (light blue). The data are the means of ± s.e.m. from 3 to 5 independent experiments.

**S5 Figure. Basolateral and apical ecto-ATPase activity of Caco-2 cells.** (A) eATP kinetics of cells exposed to eATP (0.2–7 μM). Levels of [eATP] were measured by luminometry at the basolateral (A) and apical (B) sides of the polarized Caco-2 monolayers. Data is the mean of 3 independent experiments run in duplicate. (C) Ecto-ATPase activity measured from the eATP kinetics at different [eATP] shown in panels A and B. The initial velocity of the ecto-ATPase activity was calculated by linear regression to experimental data obtaining the slope and y-intercept of the line. The slope represented the eATP hydrolysis as a function of time, *i.e*. the ecto-ATPase activity at each [eATP] and in each compartment. The points in the plot represent the mean ± s.e.m. of 3 independent experiments. The dashed lines represent a linear regression to the data allowing to obtain the ecto-ATPase kinetic constant which was 1.70 ± 0.08 and 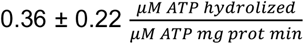 for the apical and basolateral compartments respectively.

**S6 Figure. Enzyme Vmax calculated from model fitting.** The plot shows the enzymes’ Vmax in the apical and basolateral compartments, and in non-polarized cells. The ecto-NDPK Vmax were obtained from model fitting to experimental data and are the same shown in S2 Table (for the apical and basolateral compartments) and in S1 Table (for the non-polarized cells). The ecto-AK Vmax was calculated from the model parameters using the following formula: 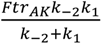, where the Ftr_AK_ was obtained from the model fitting (S2 Table for the apical and basolateral compartments and S1 Table for the non-polarized cells). The k_-2_ and k_1_ parameters value can be found in S1 Table.

**S1 Table. Mathematical model of eATP regulation in non-polarised Caco-2 cells**. Numerical values of constants were normalized by the protein cell mass in the experiments (M_cell_), measured by the Bradford method (section 2,6 in the manuscript). Parameter fitting and simulations were performed by selecting the average cell mass in the experiments (M_cell_=0.2 mg). J_L_ and J_NL_ represent the lytic and non-lytic iATP release respectively upon an osmotic shock. The value of these terms was 0 before shock application. J_leakage_ represents a constant and small iATP release observed in the absence of any stimulus. The parameter values obtained from the model fitting are expressed as the best value ± standard deviation.

**S2A Table**. JNL parameters obtained from fitting to experimental data at 180 mOsm shock in apical or basolateral compartments in polarized cells. The same value of k_obs_ was considered for both compartments. Parameters obtained from the model fitting are expressed as the best value ± standard deviation.

**S2B Table. Parameters obtained from model fitting to experimental in apical or basolateral compartments in polarized cells.** The model equations are the same shown in Table S1, however, some parameters values were fitted again to experimental data from polarised cells. The parameters whose value has change in comparison with the model of non-polarised cells are shown in this file. The rest of the parameters had the same value for non-polarised cells (shown in S1 Table). The K_ADPase_ and K_NTPase_ (for eCTP) were considered 0 in the basolateral compartment. This does not mean that there is no ecto-ADPase or ecto-NTPase activity in the basolateral side but, they can be neglected in our experimental conditions. Parameters obtained from the model fitting are expressed as the best value ± standard deviation.

